# HiCognition: a visual exploration and hypothesis testing tool for 3D genomics

**DOI:** 10.1101/2022.04.30.490134

**Authors:** Christoph C. H. Langer, Michael Mitter, Roman R. Stocsits, Daniel W. Gerlich

## Abstract

The 3D organization of the genome and epigenetic marks play important roles in gene expression, DNA repair, and chromosome segregation. Understanding how structure and composition of the chromatin fiber contribute to function requires integrated analysis of multiple genomics datasets from various techniques, experimental conditions, and cell states. Genome browsers facilitate such analysis, yet currently visualize only a few regions at a time and lack statistical functions that are often necessary to extract meaningful information. Here, we present HiCognition, a visual exploration and machine-learning tool based on a new genomic region set concept, which enables detection of patterns and associations between 3D chromosome conformation and collections of 1D genomics profiles of any type. By revealing how transcriptional activity and cohesin subunit isoforms contribute to chromosome conformation, we showcase how the flexible user interface and machine learning tools of HiCognition can help understand the relationship between structure and function of the genome.

---

Regulated expression, maintenance, and propagation of the genetic information depends not only on the DNA sequence, but also on the thousands of different proteins and posttranslational modifications that enrich at specific sites of the genome. The regulation and function of genomes further depends on an intricate organization of DNA in 3D space^1,2^, established by DNA looping^3^, chromatin phase separation^4–6^, and potentially other processes. How 3D genome organization relates to local variation in chromatin composition, DNA sequence, and physiological functions are key questions that will be important to answer for understanding the function of complex genomes. The advent of techniques mapping function, composition, and 3D organization genome-wide provides rich sources of complex data to address this challenge. Curated public repositories of various functional and 3D genomics data, e.g., Encyclopedia of DNA Elements (ENCODE)^7,8^ and 4Dnucleome^9^, provide opportunities for experimentalists to assess their data in the context of multi-dimensional epigenetic and spatial signatures. However, the challenge of extracting meaningful information from large sets of complex data has hampered progress.

A common approach towards identification of biologically relevant patterns is by studying relationships between multiple independent experiments, representing different assays, molecular components, cell states, or treatments. For example, the observation that the protein complex cohesin enriches at insulation sites of transcriptional regulation^10^ and at the boundaries of topologically associated domains (TADs)^11^ has inspired models for how the genome is organized by cohesin-mediated loop extrusion^12–14^, with broad implications for various processes^3^. Detecting associations between multiple genomics datasets is facilitated by genome browsers such as the UCSC genome browser^15–17^, which provide side-by-side views of functional genomics data and support user interaction by panning and zooming. However, available genome browsers visualize only a small number of regions at a time, which restricts the assessment of large genomes and highly heterogeneous signals in genomic profiles. To facilitate visualization and grouping of small multiples of genomic regions, a set of tools has been recently developed to leverage the concept of visual piling^18,19^. While these tools allow detection of patterns in single genomic tracks, they do not support integration of different data sources and have performance limitations with large sets of genomic views.

Systematic analysis of correlations in multiple independent genomics datasets often starts by defining a specific type of genomic region based on a common function (e.g., genes) or experimental observation (e.g., ChIP-seq peaks). Owing to the necessity to interface different datatypes and to combine algorithms from different sources, the analysis of genomic region sets is typically performed by script-based approaches^20– 22^. While script-based analysis provides flexible access to powerful statistics and machine learning tools^23–25^, it often takes a lot of time and requires advanced programming expertise to adapt workflows for investigation of new biological questions. Many wet-lab biologists have limited expertise in scripting or programming and therefore delegate advanced data analysis tasks to dedicated computer scientists, which represents a severe bottleneck in testing and developing new hypotheses.

Here, we present HiCognition, a tool for interactive visualization and statistical analysis of 3D genomics data and other (epi)genetic profiles based on a region set concept. HiCognition combines a visual exploration interface with high-performance data processing and statistical and machine learning tools. Thereby, HiCognition allows biologists without programming skills to systematically explore their large multi- dimensional genomics data, providing unprecedented opportunities for discovering fundamental mechanisms underlying the organization and function of the genome.

## Results

### Exploring genomic region sets in multi-dimensional feature space

In contrast to conventional 3D genome browsers like JuiceBox^17^ or HiGlass^16^, which visualize a specific subregion of the genome that can be panned or zoomed, HiCognition has been designed for interactive analysis of large sets of genomic regions that are pre-defined by the user before data exploration. The ***genomic region set*** approach of HiCognition allows users to address biological questions about how a specific type of region is composed, regulated, and organized in 3D space. The genomic region set can be freely defined by the user, for example, based on a common function (e.g., genes, enhancers, or origins of replication), based on molecular composition (e.g., regions with specific histone modifications or enrichment sites of proteins), or based on 3D organization (e.g., loops or topologically associated domains). The region set is provided as input data to HiCognition by a file containing genome coordinates. HiCognition then allows the user to explore associations between the genomic region set and large collections of genomics features, which can be downloaded from public repositories or from lab-internal experiments.

In HiCognition, ***genomic features*** can contain any type of numerical data associated with genomic coordinates^26–28^, including two-dimensional data like chromosome conformation contact maps (e.g., from Hi-C^29^ or SPRITE^30,31^), or one-dimensional data such as protein binding profiles (e.g., ChIP-seq^32^ or Cut&Run^33^ read densities), chromatin accessibility measurements (e. g., ATAC-seq^34^ or MNase-seq^35^), transcriptional activity (e.g., GRO-seq^36^), or replication timing measurements (e.g., Repli-seq^37^). Moreover, genomic features can contain data from unperturbed conditions as well as data obtained after genetic or chemical treatments, or data from different cell states (e.g., cell cycle stage or differentiation state), thereby enabling queries of how specific types of regions respond to perturbations or state transitions. HiCognition combines an intuitive and configurable graphical user interface with statistics and machine learning methods to enable interactive exploration of multi-dimensional genomics data within versatile workflows.

HiCognition supports data analysis by three basic approaches (Fig. 1a):

**Figure 1.**
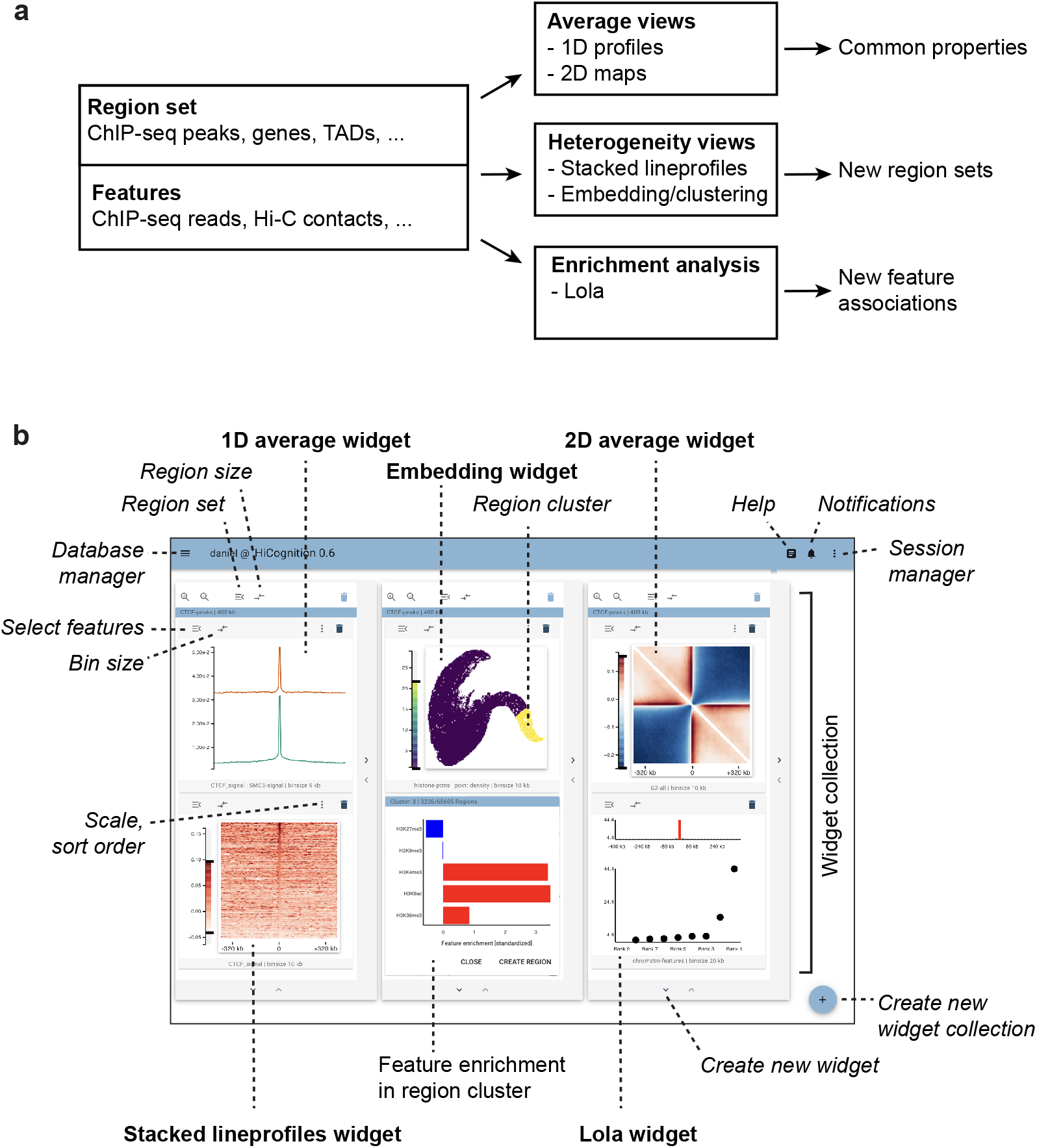
HiCognition concept and graphical user interface. **a**, Analysis workflows based on genomic region sets and collections of 3D genomics/epigenetic profiles. **b**, Graphical user interface with freely configurable widget layout. Widgets are labeled in bold and user interface elements are marked in italic. Widget collections in this figure represent visualizations of different properties of the same region set (genome-wide set of CTCF ChIP-seq peaks) with various ChIP-seq or Hi-C datasets; for explanation of individual widgets and data, please see Fig. 2 and 3 and main text.

1. ***Exploring average distributions:*** HiCognition visualizes average magnitudes of genomic signals within the region window, whereby the features can be interactively selected by the user.
2. ***Exploring region heterogeneity:*** HiCognition visualizes genomic signals of individual regions to visually explore heterogeneity in the region set. Moreover, multi-dimensional cluster analysis and visualization of region distributions in embedding plots allows identification of region sub-sets with common properties.
3. ***Enrichment analysis:*** HiCognition automatically detects features that are enriched or depleted in the specific region set under investigation relative to the genome-wide average. It further shows where within the genomic region window, individual features are particularly enriched or depleted. This enables the discovery of regulatory, functional, or spatial patterns characteristic for the region set under investigation.

The user interface of HiCognition is based on a widget architecture that allows easy configuration of views. These widgets represent genomic features and are arranged within widget collections that are associated with a specific genomic region set (Fig. 1b). This arrangement maps the abstract region set concept to a specific user interface component, allowing users to construct views that integrate different genomic features to understand the properties of a genomic region set. Specifically, following import and pre-processing of region and feature data sets, HiCognition widgets generate average feature signal plots of all regions, as well as stacked representations of individual regions, whereby the graphical user interface allows interactive adjustment of region size, resolution, look-up table, contrast, etc. For automatic detection of genomic features enriched in the region set, HiCognition provides a widget for locus overlap analysis (LOLA^38^), which is displayed as a ranked feature plot. For the analysis of heterogeneity within the region set, a clustering and embedding widget automatically groups regions based on similarity in multi-dimensional feature space and represents their distribution in embedding plots. The embedding plots are interactive and display feature patterns for individual region clusters to allow fast, interactive exploration of heterogeneity within the region set. Overall, this widget architecture with interactive visualization integrates improved versions of domain-specific tools^38^ and creatively applies state-of-the-art machine learning for embeddings^39^ and clustering.

HiCognition is implemented as a web-based tool that allows performant analysis of large datasets and interactive exploration of aggregation results. The software is open source and fully containerized, such that it can run on centralized servers or locally. An integrated database for region sets and features makes HiCognition a hub for various data types from public or private sources, whereby a session concept allows sharing of insights as fully customizable views and analysis workflows with others. A showcase server for hands on experience can be accessed at https://www.hicognition.com/app.

### Revealing common patterns in region sets

To exemplify the power of HiCognition’s region set approach, we analyzed the chromatin fiber organization around all transcriptional start sites (TSS) of protein-coding genes annotated in the human genome^40^. TSS are known to frequently contact upstream and downstream regions; at the same time, TSS insulate against contacts between upstream and downstream genomic regions^41–45^. Using published ChIP-seq data from HeLa cells^8,46^, we first visualized the distribution of two key architectural regulators, cohesin (based on its subunit Structural Maintenance of Chromosomes 3, SMC3) and CCCTC-binding factor (CTCF) using HiCognition’s *1D average widget*. A prominent enrichment of both proteins at TSS (Fig. 2a, panel i) supports a role of cohesin-mediated DNA looping in shaping the conformation around TSS^10,41,47,48^.

**Figure 2.**
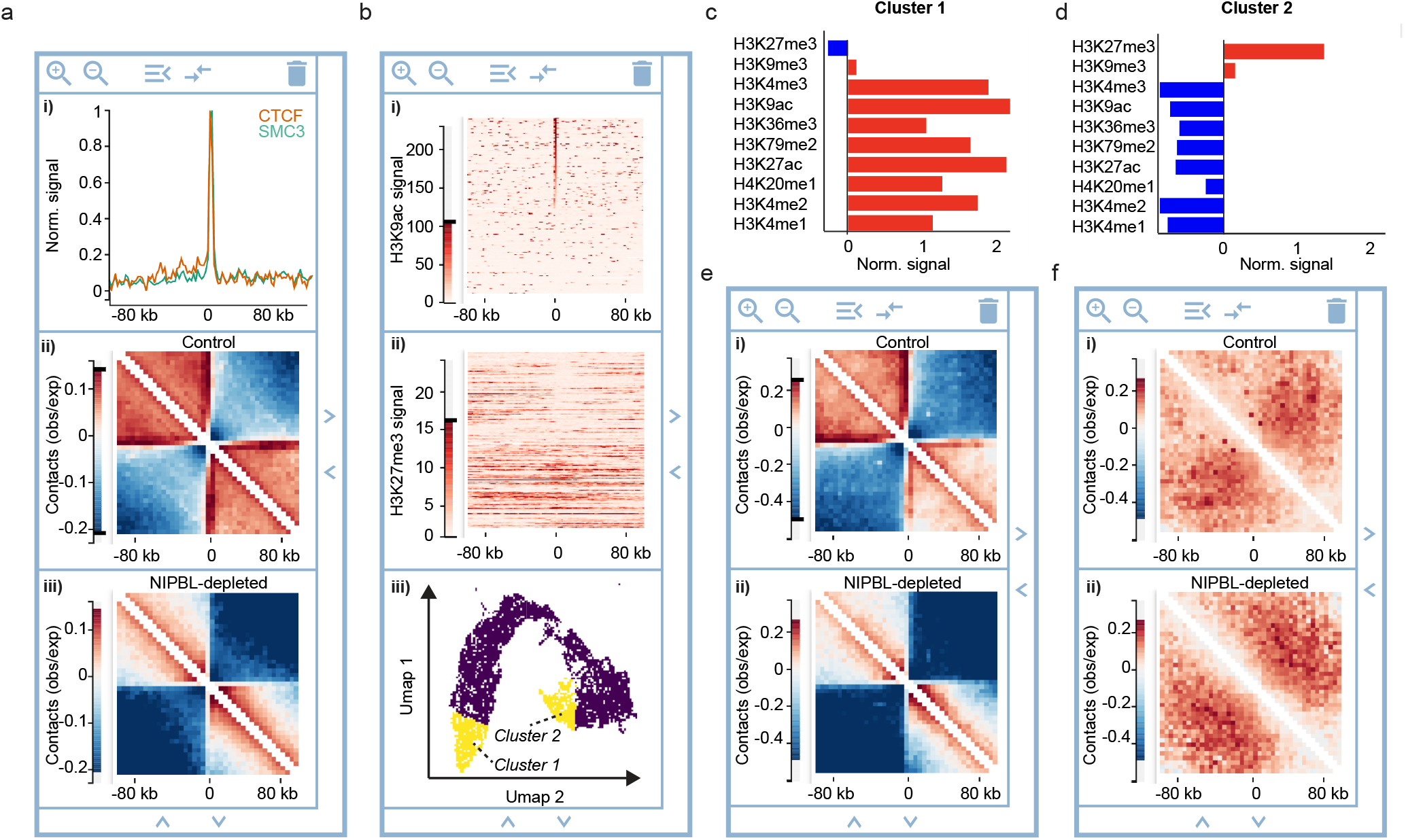
Exploring average profiles and heterogeneity in region sets. A genome-wide region set of protein-coding genes was analyzed based on published ChIP-seq and Hi-C data, centered on the TSS. Data show averages of genes localized on the forward strand, with the gene body facing towards the right from the TSS (center coordinate). **a**, Average ChIP-seq reads of CTCF and SMC3 (i), average Hi-C contact maps of unperturbed wildtype cells (ii), and average Hi-C contact maps of NIPBL-depleted cells (iii). b, Heterogeneity of histone posttranslational modifications within the protein-coding gene region set, visualized by stacked line profiles sorted by the read density of H3Kac (i) and displayed for H3K27me3 ChIP-seq read density (ii), with the sorting order coupled to i, and regional heterogeneity analysis based on 10 histone posttranslational modifications by embedding and clustering (iii). **c, d**, Regional subsets created by clustering as shown in b (iii) were analyzed for 10 different histone posttranslational modifications. Red indicates enrichment, blue indicates depletion. **c**, Cluster 1 contains TSS regions of protein-coding genes enriched in marks for actively transcribed chromatin. **d**, Cluster 2 contains TSS regions enriched in marks for transcriptionally repressed chromatin. **e, f**, Chromosome conformation analysis around TSS region subsets as shown in c, d. **e**, HiC average contact maps of Cluster 1 from unperturbed wildtype cells (i) and cells depleted of NIPBL (ii). **f**, HiC average contact maps as in (e) for Cluster 2.

To assess the 3D organization of protein-coding genes, we next visualized the genome-wide average contact probability around TSS using the *2D average widge*t and published Hi-C data^49^ (Fig. 2a, panel ii). Prominent stripes emerging from the TSS towards upstream and downstream regions indicate frequent interactions of TSS with distal genomic regions. Moreover, contacts within regions upstream or downstream the TSS were much more frequent than between upstream and downstream regions (Fig. 2a, visible as red and blue areas, respectively), as previously observed^41–44^. Thus, HiCognition allows simple visualization of genome-wide averages for region-type-specific conformations.

To assess the functional role of cohesin-mediated looping to the conformation at TSS, we next used the *2D average widget* to visualize published Hi-C data obtained from cells depleted of Nipped-B-like protein (NIPBL)^49^, a cofactor essential for cohesin-mediated loop extrusion^50,51^ (Fig. 2a, panel iii). The stripes emerging from TSS and the squared regions containing high contact probability that were characteristic for unperturbed controls were almost completely suppressed in the Hi-C maps obtained from NIPBL-depleted cells, indicating a key role of cohesin-mediated looping in establishing these structures, consistent with previous observations^41,48^. Thus, HiCognition enables fast and interactive side-by-side visualization of genome-wide average profiles across various techniques and experimental conditions.

### Understanding heterogeneity within region sets

Understanding the relationship between chromatin fiber composition, 3D conformation, and physiological function has remained challenging owing to the heterogeneity of regions defined by a common feature under investigation. HiCognition’s region set approach allows fast and simple visualization of regional heterogeneity and supports interactive clustering of these regions based on multiple genomic features. To demonstrate how HiCognition’s flexible widget architecture can be used for heterogeneity analysis of region sets, we investigated how histone posttranslational modification patterns relate to chromosome conformation around genes. Using the *Stacked lineprofiles widget*, we visualized for the genome-wide set of TSS regions the ChIP-seq read densities of two histone posttranslational modifications, H3K9ac and H3K27me3, which enrich at transcriptionally active or inactive chromatin, respectively^52,53^. Sorting the line profiles by H3K9ac abundance showed that only about half of the TSS regions were enriched for this mark (Fig. 2b, panel i). Moreover, displaying stacked line profiles of H3K27me3 ChIP-seq read density in a separate widget and sharing the sort order between widgets showed that TSS regions enriched in H3K9ac are depleted of H3K27me3 (Fig. 2b, panel ii). Thus, coupling multiple widgets by sorting allows intuitive visual assessment of correlations between genomic features.

Next, we aimed to identify region subsets with distinct histone modification profiles for the study of the corresponding Hi-C conformations, considering an extended set of ten different histone posttranslational modifications (see methods for details). HiCognition’s *Embedding widget* visualizes regional heterogeneity based on multi-dimensional feature values, which can contain linear profiles such as ChIP-seq data or Hi-C contact matrices (Fig. 2b, panel iii). Besides visualizing heterogeneity, the *Embedding widget* automatically groups regions into clusters by feature similarity. The features enriched or depleted in each cluster are interactively displayed by pointing to clusters. We selected two clusters enriched either in marks for transcriptionally active chromatin or transcriptionally repressed chromatin (Fig. 2b-d) to create two new region subsets for analysis of the corresponding Hi-C conformations.

Using the *2D average widget* and the Hi-C data of HeLa cells, we observed pronounced high-contact stripes and insulation around TSS for the region subset enriched in active chromatin marks, whereas these Hi-C structural features were entirely absent in the region subset enriched in repressive histone marks (Fig. 2e, f, panels i), consistent with previous script-based analyses of mouse stem cell data^41^. To investigate how cohesin-mediated DNA looping contributes to chromosome conformation at TSS residing in transcriptionally active or inactive chromatin, we visualized average Hi-C maps of NIPBL-depleted cells, using published data^49^. For the region subset enriched in transcriptionally active histone marks, we found strong reduction of stripes and insulation around TSS, whereas the region subset with repressive marks was unaffected by NIPBL depletion (Fig. 2e, f, panels ii). Together, these data suggest that cohesin-mediated DNA looping establishes a specific chromosome architecture around transcriptionally active TSS but not at inactive TSS. Thus, HiCognition’s flexible widget architecture enables simple and powerful analysis workflows to explore regional heterogeneity and to detect interactions between different types of genomics data.

### Discovering new associations with HiCognition

Public repositories such as ENCODE^8^ or the 4Dnucleome^9^ contain thousands of different genomics data sets derived from diverse technologies, cell types, and experimental conditions. The difficulty to interpret such complex data has prompted the development of various computational methods to detect associations between specific types of regions and features describing the chromatin fiber, such as GREAT^54^, the Encode ChIP-seq significance tool^55^, GenometriCorr^56^ and Locus Overlap Analysis (LOLA)^38^. HiCognition provides an improved implementation of LOLA, extended by interactive exploration of feature enrichment in distinct genomic sub-bins obtained from a region set. We exemplify association analysis with HiCognition’s *Lola widget* by investigating how cohesin subunit isoforms relate to chromosome conformation.

Cohesin contains three core subunits that form a ring, and an associated Stromal Antigen (STAG) subunit of which vertebrates encode two isoforms, STAG1 and STAG2^57–60^. Previous script- based analysis of ChIP-seq profiles and Hi-C data showed that STAG2-cohesin predominantly forms loops at active TSS, whereas STAG1-cohesin predominantly contributes to the formation of TADs^58,61–63^. Here, we aim to recapitulate these findings and search for new associations by the automated machine learning tools and interactive workflows of HiCognition. We created a region set centered on all 34,857 SMC3 ChIP-seq peaks and then clustered SMC3 regions based on the abundance of STAG1 and STAG2, using the *Embedding widget* and published ChIP-seq data^63^ (Fig. 3a, b). Comparing ChIP-seq read densities with the *1D average widget* showed that the region subset enriched in STAG1 contained less SMC3 than the region subset enriched in STAG2 (Fig. 3c, d).

**Figure 3.**
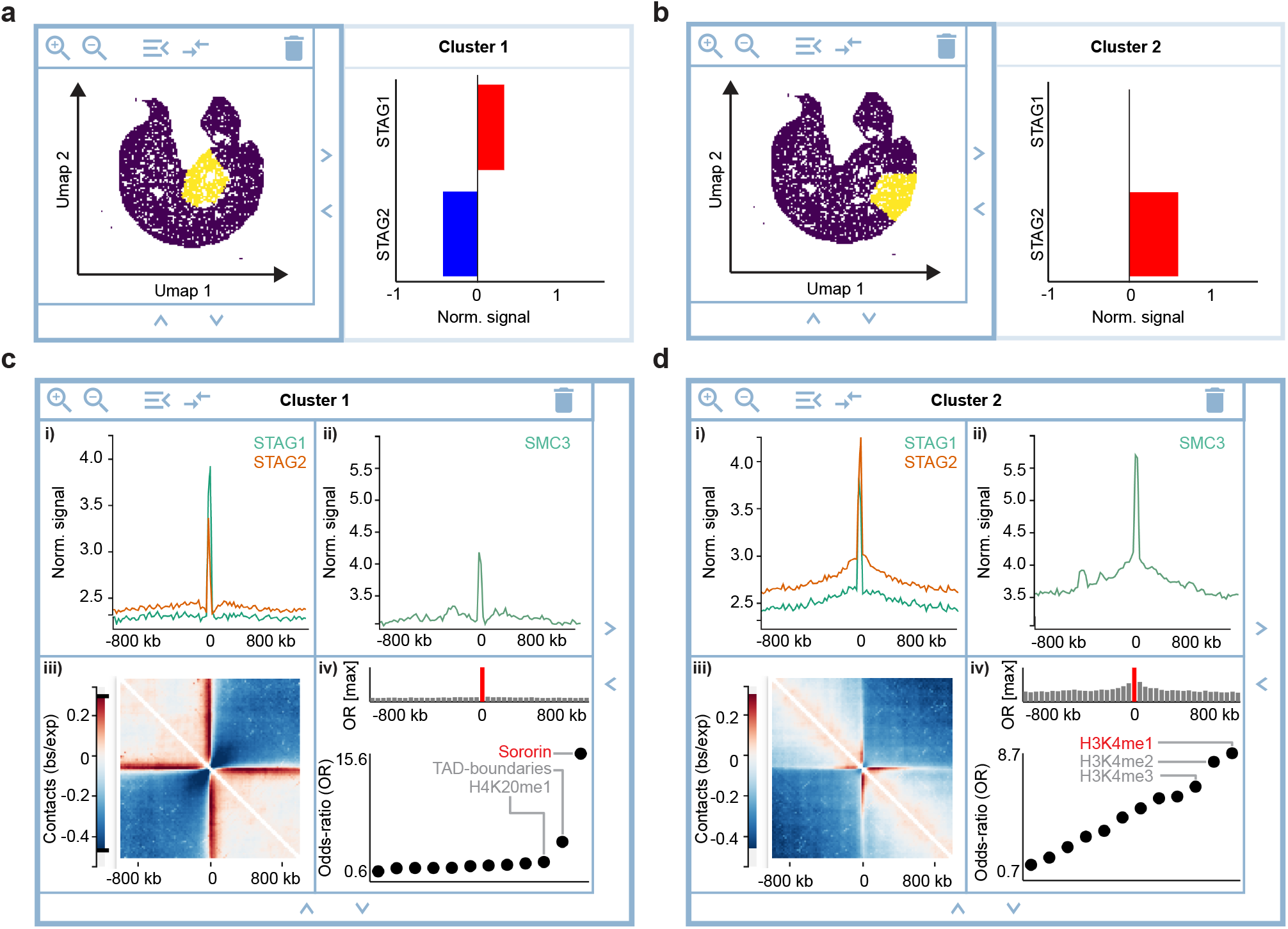
Detecting associations with HiCognition. **a, b**, Visualization of regional heterogeneity and clustering with the *Embedding widget* for 34,857 SMC3 ChIP-seq peaks based on the ChIP-seq read densities of STAG1 and STAG2. **a**, Cluster 1, representing SMC3 ChIP-seq peaks enriched in STAG1. **b**, Cluster 2, representing SMC3 ChIP-seq peaks enriched in STAG2 ChIP-seq reads. **c, d**, analysis of common patterns and associations for Cluster 1 and 2 as in a, b. **c**, Average read density of STAG1 and STAG2 (i) and SMC3 (ii), average Hi-C contact map (iii), and LOLA analysis for associations with 11 region data sets. **d**, analysis as in c for Cluster 2.

To visualize the chromosome conformation around these region subsets, we used the *2D average widget* and published Hi-C data^49^. Strikingly, the STAG1-enriched sites had much more pronounced long-range contacts than the STAG2-enriched sites (Fig. 3c, d, panels iii), despite the lower abundance of the core cohesin subunit SMC3 at STAG1-enriched sites (Fig. 3c, d, panels ii). To determine in which genomic context STAG1- or STAG2-enriched sites predominantly reside, we used the Lola widget to analyze 11 region sets including histone posttranslational modifications, TAD boundaries, and the cohesin-associated protein Sororin that is required for cohesion maintenance in G2^64,65^. This analysis showed that STAG1- enriched sites predominantly reside at TAD boundaries, whereas STAG2-enriched SMC3 peaks predominantly reside in chromatin bearing marks of active transcription (Fig. 3c, d, panels iv), supporting the previously reported distinct localization and function of cohesin bound to STAG1 or STAG2, respectively^58,61–63^. Moreover, STAG1-enriched cohesin sites also prominently overlapped with Sororin sites detected by ChIP-seq in G2 phase of the cell cycle^46^, indicating a previously unrecognized association between genomic sites of sister chromatid cohesion and genomic sites where STAG1-enriched cohesin forms long-range loops in G1. Importantly, HiCognition’s region-set-based approach and flexible widget architecture enable detection of such complex associations within a few minutes. Thus, HiCognition allows biologists untrained in genomic analysis to rapidly perform their own analyses, discover new associations, and generate new hypotheses, greatly reducing the bottleneck between data generation and interpretation.

## Discussion

HiCognition leverages interactive genome exploration to comprehensive views of genome-wide region sets defined by a common property. Its flexible user interface and integrated statistics and machine learning tools support the detection of common patterns, heterogeneity, and associations in complex genomics datasets representing 3D conformation, epigenetic profiles, and functional readouts. A fast and computationally efficient implementation allows real-time browsing through thousands of genomic regions, thereby accelerating hypothesis testing on genomics data of various experimental techniques, experimental conditions, or cell states.

HiCognition’s rich online documentation and containerized distribution supporting desktop as well as server installations provide easy access for both experienced developers as well as beginner analysts. The integrated database and interfaces to widely used file formats allow assessment of a biologist’s own data in the context of the vast amount of public data available from resources like ENCODE or 4Dnucleome. HiCognition’s streamlined workflows and visualization concepts enable users to address a broad range of biological questions, yet the focus on usability limits customizability compared to approaches that simply provide a graphical interface to command-line tools^66^ or custom scripts^67^. Via the export of region set coordinates derived from clustering and association analysis, however, HiCognition can be seamlessly integrated with script-based analysis for extended functionality. Hence, HiCognition allows biologists lacking programming skills to rapidly reduce the space of possible hypotheses before applying more time-consuming methods. Furthermore, the software’s modular design and open-source implementation in Python provide an extendable framework towards development of new machine learning algorithms and visualization concepts. Therefore, we foresee that HiCognition will serve as a bridge between the experimentalists who formulate biological hypotheses and specialized computer scientists implementing script-based analyses workflows.

While HiCognition’s potential is exemplified here by an analysis of epigenetic marks and topological structures formed by cohesin, the software is applicable to any type of 1D or 2D genomics data. Its ease of use and data integration based on the region set concept will provide new opportunities for discovering relationships between structure and function of the genome.

## Acknowledgments

The authors thank Jan-Michael Peters, Paul D. Batty, Federico Teloni, Zsuzsanna Takacs, and Sofia Kolesnikova for comments on the manuscript. This project has received funding from the European Research Council (ERC) under the European Union’s Horizon 2020 research and innovation program (grant agreement No 101019039), from the Austrian Academy of Sciences, and the Vienna Science and Technology Fund (WWTF; project nr. LS17-003).

## Author contributions

Conception: M.M., C.C.H.L, D.W.G.; software design and implementation: M.M., C.C.H.L.; data analysis and interpretation: D.W.G., M.M., R.R.S.; manuscript writing: M.M., C.C.H.L, D.W.G.; funding acquisition and supervision: D.W.G.

## Methods

### Software architecture

HiCognition is a containerized application (https://github.com/docker/compose) and designed as a server-client web app to minimize set-up requirements and facilitate easy usage for non- technical users after set-up (Fig. S1a).

The backend portion of HiCognition is implemented as a Flask webserver (https://github.com/pallets/flask) with NGINX (https://github.com/nginx) as a reverse proxy that operates in conjunction with a MySQL database (https://github.com/mysql) to persist metadata and data preprocessing results. The server utilizes a Redis task queue (https://github.com/rq/rq) to offload time-intensive computation tasks to an adjustable number of worker containers. The communication between these workers and the main server is implemented via network requests (when submitting a task) and the MySQL database (when registering a task as complete). This organization allows the operation of the worker containers on separate machines that could, in principle, be started on demand.

The frontend part of HiCognition is implemented in JavaScript and uses the Vue.js framework (https://github.com/vuejs/vue) to manage components and implement reactivity. The visualizations are custom-designed for each type of data widget (see below for details) and are implemented either using the data-driven visualization library D3.js (https://github.com/d3/d3) or in case of more demanding visualizations using PixiJS (https://github.com/pixijs/pixijs).

For implementation details of the HiCognition architecture, see the GitHub repository (https://github.com/gerlichlab/hicognition) and the accompanying documentation page (https://gerlichlab.github.io/hicognition/docs/).

### Point- and interval-regions

As genomic data frequently span multiple length-scales ^16,17^, visualization concepts have to adapt to this challenge. HiCognition solves this problem by precomputing a “resolution- stack” for each genomic region-set (Fig. S1b). This precomputation is adapted for two types of genomic regions supported by HiCognition:

*Point-regions* are specified by center coordinates and the region surrounding the center position can be adjusted interactively for analysis and visualization. This enables the user to zoom in and out of genomic regions when viewing data to discover genomic effects at multiple length scales.

*Interval-regions* are specified by start and end coordinates and each region includes 20% neighboring regions on either side. The processing bin size for this region type is adjusted by normalization to the interval size, and thus different for differently sized regions. Interval regions allow to investigate length-independent patterns, as for example profiles of genes that are scaled to transcription start and termination sites.

### Data management and preprocessing

HiCognition contains a dataset manager that stores available datasets as well as finished pre-computations in a MySQL database. The user interface of HiCognition distinguishes between two principal types of data – genomic regions of interest and genomic features that are available for precomputation (Fig. S2a). Users can add and view datasets in an interactive table that allows filtering and editing (Fig. S2b).

HiCognition supports the most common input data formats for genomic regions and features. Specifically, genomic regions can be added as bed-files^15^, 1D-features as bigwig files^68^ and 2D-features as cooler files^20^. These files can be uploaded one at a time or using a bulk upload feature (see our documentation at https://gerlichlab.github.io/hicognition/docs/data_management/ for details).

To select a region-set of interest, the user can submit preprocessing tasks using the preprocessing dialogues and get an overview of running and finished computations via the dataset viewer of the genomic regions (Fig. S2c). Once pre-computation of a combination of a region-set of interest and a genomic feature has finished, it is available for display.

Many preprocessing steps involve analysis of genomic feature collections, for example, when calculating enrichment amongst a set of candidate features or embedding regions based on the values of multiple features (see below for details). In HiCognition, users can create feature collections in a specific dialogue window and select them for preprocessing and display.

HiCognition also supports adding and managing multiple genome assemblies to analyze and compare data generated for different genome assemblies and species.

### Data and workflow sharing

HiCognition’s allows storing specific arrangements of widgets, widget collections, and the corresponding data under display as named sessions. This is possible due to an implementation of the HiCognition analysis view as declarative configurations stored in the Vuex frontend storage (https://github.com/vuejs/vuex/). Here, the arrangement, settings, and data sources loaded in a particular widget are stored as JavaScript objects, and HiCognition reacts to changes therein by adjusting the displayed view. This makes it easy to restore saved sessions from configuration objects stored in the database and to share saved sessions with collaborators through a static link.

### Widgets and visualization concepts

HiCognition uses widget-collections as a container to display specific visualizations (Fig. 1b). A widget collection has a single region-set that is shared by all its contained widgets. Each widget in the collection represents a genomic feature or a collection of genomic features and provides a suitable visualization for the respective data (Fig. 1b).

#### 1D-average widget

The 1D-average widget displays the average magnitude of a 1D genomic feature, as for example ChIP-seq reads, for the selected region set in the widget collection as a line plot. The preprocessing algorithm extracts snippets of the relevant genomic feature for each genomic region and calculates the average value over all snippets along the relative genomic offset.

#### 2D-average widget

The 2D-average widget displays the average magnitude of a 2D- genomic feature, for example a Hi-C contact probability map, for the selected region set in the widget collection as a 2D heatmap. The preprocessing algorithm extracts snippets of the 2D- genomic feature for each rectangular genomic region and calculates the average value over all snippets for each pixel.

#### Stacked line profile widget

The stacked line profile widget displays individual examples of 1D-genomic features for the selected region set in the widget collection as a 2D heatmap. Within this heatmap, each row represents a specific genomic region. The preprocessing algorithm extracts the relevant genomic feature snippets for each genomic region (subsampled to contain a maximum of 1000 regions) and “stacks” them vertically to form a matrix for display.

#### 1D-feature embedding widget

The 1D-feature embedding widget displays the distribution of genomic regions based on a collection of 1D genomic features. The results are displayed as a 2D-histogram, where points close on the plot represent genomic regions with similar feature profiles. The dimensionality reduction algorithm UMAP^39^ is used with default parameters to embed the high-dimensional regions into a two-dimensional space suitable for display.

This widget also automatically groups region neighborhoods by k-means clustering, with 10 (“large neighborhood”) or 20 (“small neighborhood”) clusters, respectively, in the embedded space. The normalized intensity of the features for each cluster is then calculated and used to interactively display the distribution of features within the selected clusters by mouse hovering. Users can create new regions from interesting subsets by clicking on a subset and giving it a name in the relevant dialogue.

#### 2D-feature embedding widget

The 2D-feature embedding widget displays the distribution of genomic regions using a single 2D genomic feature. The results are displayed as a 2D-histogram, where points next to each other exemplify genomic regions with similar 2D-feature values. The widget implements a hover interaction that shows the 2D average with respect to the selected genomic feature for the selected subset. Users can create new regions from interesting subsets by clicking on a subset and giving it a name in the relevant dialogue.

The preprocessing algorithm extracts snippets of the 2D genomic feature for each genomic region in the region set. These snippets are then smoothed using a Gaussian filter and down- sampled to be of size 10 × 10. Here, the smoothing kernel size and standard deviation depend on the interpolation factor:

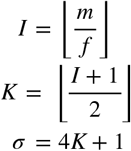

Where *I* is the interpolation factor, *m* is the size of the quadratic snippet, *f* is the target size of the down-sampled matrix (in this case 10), *K* is the size of the smoothing kernel, and *σ* is the standard deviation of the Gaussian filter. The smoothing and down-sampling operations are done using OpenCV (https://github.com/opencv/opencv). Note that since the snippets can be of different sizes (see above for details), the interpolation factor and smoothing function can differ for different extracted snippets. The down-sampled matrix is then flattened and treated as image features for each of the genomic regions, resulting in a matrix where each row corresponds to a genomic region in the region set and each column to one of the pixel features (100 in total). Then, the matrix is embedded into a 2D space using UMAP^39^ (https://github.com/lmcinnes/umap), and clustering is performed as for the 1D-feature embedding widget. The representation for each cluster that is displayed to the user is the 2D average of all contained matrix snippets in the original pixel space.

#### Association widget

The association widget allows users to quantify for a given region set the extent by which other sets of independent genomic regions overlap, based on the LOLA method^38^. This allows to detect associations between different types of genomics data, as for example ChIP-seq peaks and Hi-C structures like boundaries of TADs.

This widget consists of two visualizations, where the upper bar chart shows quantification of the maximum enrichment of all regions within a collection, and the lower chart indicates the enrichment values for a selected bar ranked by enrichment. A significantly faster python reimplementation of LOLA^38^ (https://github.com/Mittmich/pylola) allows calculating the association not just on the region of interest level but for each individual bin of these regions. Specifically, we use a bin as the target region, the regions in the selected collection as query regions, and all genomic-wide bins of that size as a universe. The reported values correspond to the odds ratio of the underlying contingency table for each combination of target, query, and universe.

### Use-cases

#### Data sources

All data sets used for analysis in the current study have been obtained from public repositories as listed in the following table:

| Name | Repository | ID | Reference |
| --- | --- | --- | --- |
| SMC3 Chip-Seq reads | ENA | SAMEA5988740 and SAMEA5988741 | Ladurner et al.<sup>46</sup> |
| Sororin Chip-Seq reads | ENA | SAMEA3716450 and SAMEA3716449 | Ladurner et al.<sup>46</sup> |
| CTCF read density | GEO | GSM733785 | Encode Consortium<sup>8</sup> |
| G2 Hi-C data WT | GEO | GSM4613674 | Mitter et al.<sup>49</sup> |
| G2 Hi-C data NIPBL depleted | GEO | GSM4613678 | Mitter et al.<sup>49</sup> |
| Stag1 Chip-Seq read density | GEO | GSM4106803 | Wutz et al.<sup>63</sup> |
| Stag2 Chip-Seq read density | GEO | GSM4106804 | Wutz et al.<sup>63</sup> |
| H3K4me1 Chip-Seq peaks | GEO | GSM798322 | Encode Consortium<sup>8</sup> |
| H3K4me2 Chip-Seq peaks | GEO | GSM733734 | Encode Consortium<sup>8</sup> |
| H3K4me3 Chip-Seq peaks | GEO | GSM733682 | Encode Consortium<sup>8</sup> |
| H3K9me3 Chip-Seq peaks | GEO | GSM1003480 | Encode Consortium<sup>8</sup> |
| H3K9ac Chip-Seq peaks | GEO | GSM733756 | Encode Consortium<sup>8</sup> |
| H3K79me2 Chip-Seq peaks | GEO | GSM733669 | Encode Consortium<sup>8</sup> |
| H3K27ac Chip-Seq peaks | GEO | GSM733684 | Encode Consortium<sup>8</sup> |
| H4K20me1 Chip-Seq peaks | GEO | GSM733689 | Encode Consortium<sup>8</sup> |
| H3K36me3 Chip-Seq peaks | GEO | GSM733711 | Encode Consortium<sup>8</sup> |
| H3K27me3 Chip-Seq peaks | GEO | GSM733696 | Encode Consortium<sup>8</sup> |
| H3K27me3 Chip-Seq read density | GEO | GSM733696 | Encode Consortium<sup>8</sup> |
| H3K9me3 Chip-Seq read density | GEO | GSM1003480 | Encode Consortium<sup>8</sup> |
| H3K4me3 Chip-Seq read density | GEO | GSM733682 | Encode Consortium<sup>8</sup> |
| H3K9ac Chip-Seq read density | GEO | GSM733756 | Encode Consortium<sup>8</sup> |
| H3K36me3 Chip-Seq read density | GEO | GSM733711 | Encode Consortium<sup>8</sup> |
| H3K79me2 Chip-Seq read density | GEO | GSM733669 | Encode Consortium<sup>8</sup> |
| H3K27ac Chip-Seq read density | GEO | GSM733684 | Encode Consortium<sup>8</sup> |
| H4K20me1 Chip-Seq read density | GEO | GSM733689 | Encode Consortium<sup>8</sup> |
| H3K4me2 Chip-Seq read density | GEO | GSM733734 | Encode Consortium<sup>8</sup> |
| H3K4me1 Chip-Seq read density | GEO | GSM798322 | Encode Consortium<sup>8</sup> |
| gencode.v38lift37.basic.annotation.gtf | GENCODE | Release38 GRCh37 | Gencode Consortium<sup>40</sup> |
| G2 Hi-C data WT TADs | Github | github.com/gerlichlab/scshic_analysis/blob/master/data/TADs_final.bedpe | Mitter et al.<sup>49</sup> |

#### Preparation of datasets for HiCognition

All ChIP-seq data were directly imported into HiCognition based on data from public repositories, except for the SMC3 and Sororin ChIP-seq peaks, which were detected by the following procedure in the published ChIP-seq read profiles from Ladurner et al.^46^:

Deep (Illumina) sequencing results of ChIP-Seq libraries were downloaded from ENA (ID: SAMEA5988740) and mapped against the human hg19 reference assembly using bowtie resp. bowtie2 (http://bowtie-bio.sourceforge.net/bowtie2/index.shtml) counting only uniquely mappable reads with 0 - 2 mismatches allowed. Resulting alignments from two replicates each were processed with MACS peak calling algorithm (version 1.4.2) with a P-value threshold of 1e-10 resp. 1e-5 adding control inputs from the same cell line. Peak overlaps were calculated by using multovl 1.3 (https://github.com/aaszodi/multovl) while treating overlaps as unions and including unique peaks from both replicates. Since occasionally two neighboring peaks from one dataset overlap with a single peak in another dataset, the output of such overlap is displayed as a connected genomic site and merged into one single data entry.

To derive protein-coding genes split along their direction of transcription, the GENCODE annotations for hg19 (GRCh37) were downloaded and filtered for entries that were of type “gene” and of gene type “protein_coding”. These genes were then split into genes with strand “+”, named “forward”, and genes with strand “-”, named “reverse”. The transcriptional start sites for these genes were then defined to be the start or end of these intervals respectively and saved as bed files. The script for this preprocessing step can be found in the HiCognition GitHub repository (https://github.com/gerlichlab/hicognition/blob/master/publication/scripts/ convert_genes.ipynb). For the use-case figures, the transcriptional start sites of “forward” oriented genes were used.

### Showcase server

To provide readers a fast hands-on experience of HiCognition, we implemented a showcase server (www.hicognition.com/ app). On this server, the login for individual users is deactivated. We uploaded and preprocessed all the datasets in this paper so the reader can explore them independently and provide all saved sessions used for the figures in this paper. On this server, the upload and preprocessing functionality is deactivated.

**Supplementary Figure 1.**
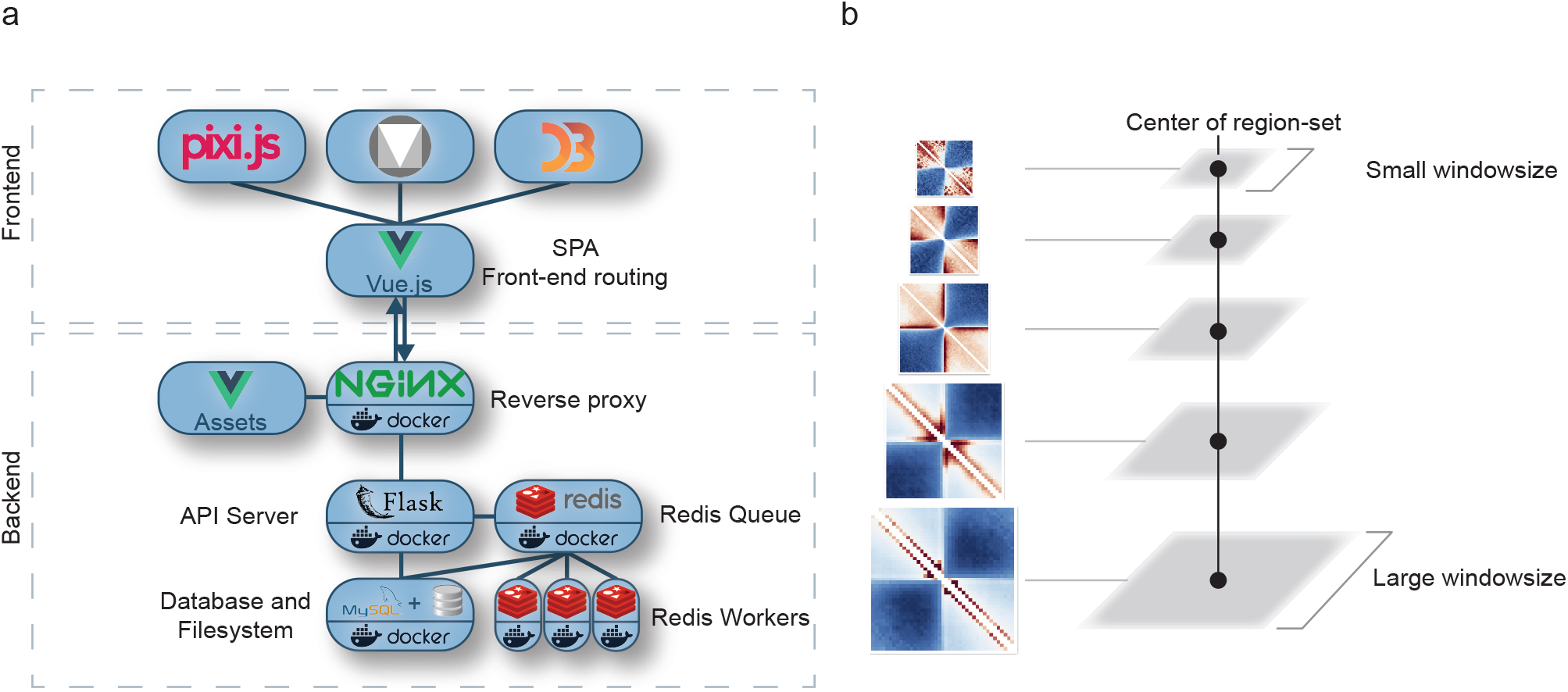
Implementation of HiCognition. **a**, HiCognition is a single-page application connected to an API backend server. The backend components are orchestrated and containerized via docker and consist of a NGINX web server, the Flask API server that dispatches high-performance precomputation tasks into a Redis queue, and a MySQL database for persistence. The frontend is based on the Vue.js JavaScript framework. The user interface is built with “Vue Material” components. The custom visualizations are built with the d3.js visualization library and the pixi.js rendering library. **b**, HiCognition precomputes a resolution stack with different window sizes and resolutions around a genomic region set to allow real-time exploration of its multi-scale neighborhood.

**Supplementary Figure 2.**
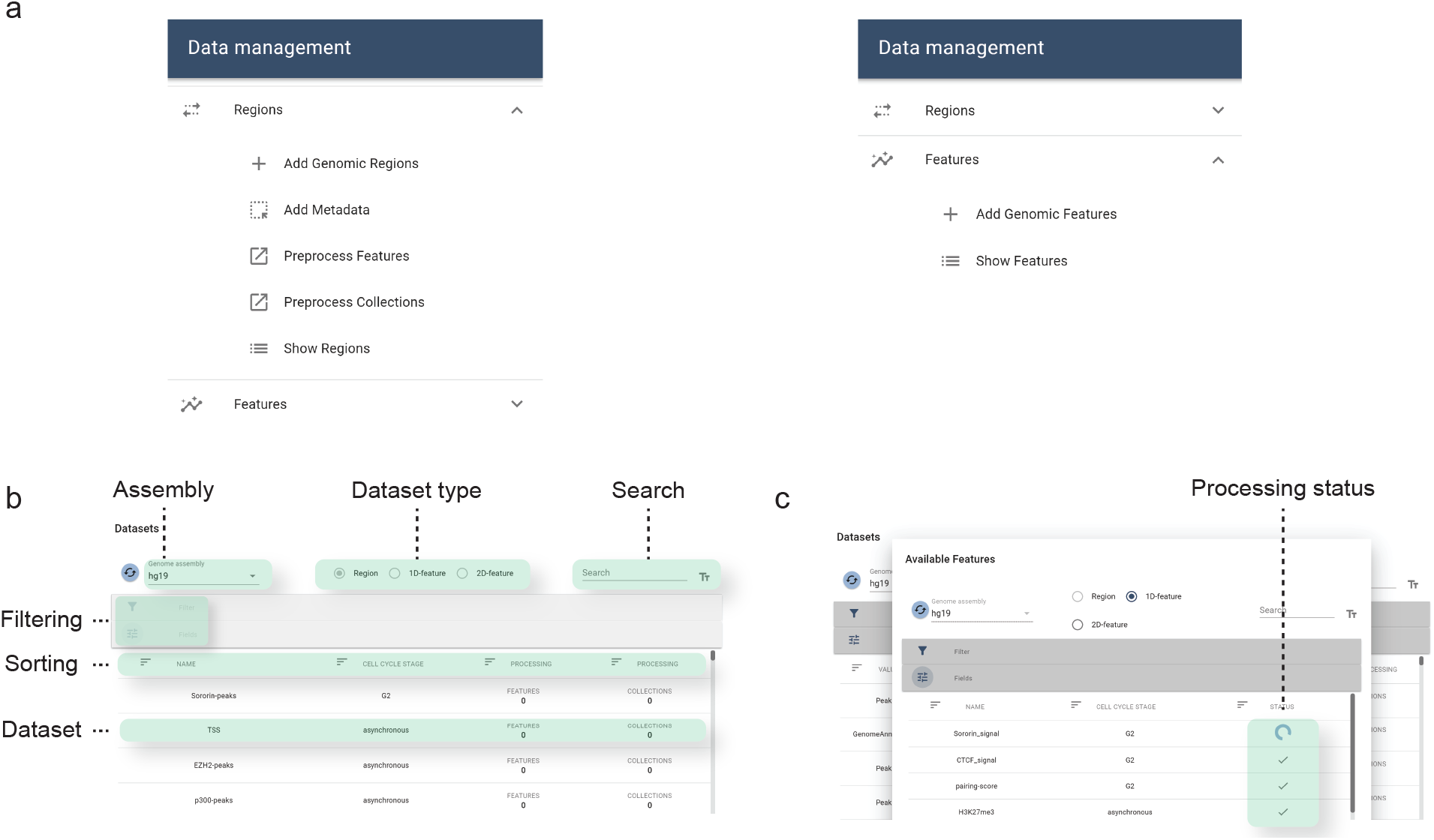
User interface for data set management. **a**, HiCognition provides conceptual separation between genomic regions of interest and genomic features. This is captured in the data management functionality by separating the options to upload, preprocess and edit genomic regions and features. **b**, HiCognition provides an interactive dataset table for managing genomic datasets. This includes selecting genome assemblies, filtering on metadata, searching for datasets, and modifying and deleting datasets. **c**, Within the dataset table, the processing state of genomic features for a specific genomic region set can be viewed within a processing dialogue.

## References

1. Misteli, T. The Self-Organizing Genome: Principles of Genome Architecture and Function. Cell 183, 28–45 (2020).

2. Dekker, J. & Mirny, L. The 3D Genome as Moderator of Chromosomal Communication. Cell 164, 1110–1121 (2016).

3. Davidson, I. F. & Peters, J.-M. Genome folding through loop extrusion by SMC complexes. Nat. Rev. Mol. Cell Biol. 22, 445–464 (2021).

4. Gibson, B. A. et al. Organization of Chromatin by Intrinsic and Regulated Phase Separation. Cell 179, 470–484.e21 (2019).

5. Erdel, F. & Rippe, K. Formation of Chromatin Subcompartments by Phase Separation. Biophys. J. 114, 2262–2270 (2018).

6. Mirny, L. A., Imakaev, M. & Abdennur, N. Two major mechanisms of chromosome organization. Curr. Opin. Cell Biol. 58, 142–152 (2019).

7. ENCODE Project Consortium et al. Expanded encyclopaedias of DNA elements in the human and mouse genomes. Nature 583, 699–710 (2020).

8. Consortium, T. E. P. An integrated encyclopedia of DNA elements in the human genome. Nature 489, 57–74 (2012).

9. Dekker, J. et al. The 4D nucleome project. Nature 549, 219–226 (2017).

10. Wendt, K. S. et al. Cohesin mediates transcriptional insulation by CCCTC-binding factor. Nature 451, 796–801 (2008).

11. Rao, S. S. P. et al. A 3D map of the human genome at kilobase resolution reveals principles of chromatin looping. Cell 159, 1665–1680 (2014).

12. Alipour, E. & Marko, J. F. Self-organization of domain structures by DNA-loop-extruding enzymes. Nucleic Acids Res. 40, 11202–11212 (2012).

13. Fudenberg, G. et al. Formation of Chromosomal Domains by Loop Extrusion. Cell Rep. 15, 2038–2049 (2016).

14. Sanborn, A. L. et al. Chromatin extrusion explains key features of loop and domain formation in wild-type and engineered genomes. Proc. Natl. Acad. Sci. U. S. A. 112, E6456–65 (2015).

15. Kent, W. J. et al. The human genome browser at UCSC. Genome Res. 12, 996–1006 (2002).

16. Kerpedjiev, P. et al. HiGlass: web-based visual exploration and analysis of genome interaction maps. Genome Biol. 19, 125 (2018).

17. Durand, N. C. et al. Juicebox Provides a Visualization System for Hi-C Contact Maps with Unlimited Zoom. Cell Syst 3, 99–101 (2016).

18. Lekschas, F., Bach, B., Kerpedjiev, P., Gehlenborg, N. & Pfister, H. HiPiler: Visual Exploration of Large Genome Interaction Matrices with Interactive Small Multiples. IEEE Trans. Vis. Comput. Graph. 24, 522–531 (2018).

19. Lekschas, F. et al. A Generic Framework and Library for Exploration of Small Multiples through Interactive Piling. IEEE Trans. Vis. Comput. Graph. 27, 358–368 (2021).

20. Abdennur, N. & Mirny, L. A. Cooler: scalable storage for Hi-C data and other genomically labeled arrays. Bioinformatics 36, 311–316 (2020).

21. Ramírez, F. et al. High-resolution TADs reveal DNA sequences underlying genome organization in flies. Nat. Commun. 9, (2018).

22. Gentleman, R. C. et al. Bioconductor: open software development for computational biology and bioinformatics. Genome Biol. 5, R80 (2004).

23. Harris, C. R. et al. Array programming with NumPy. Nature 585, 357–362 (2020).

24. Virtanen, P. et al. SciPy 1.0: fundamental algorithms for scientific computing in Python. Nat. Methods 17, 261–272 (2020).

25. Pedregosa, F. et al. Scikit-learn: Machine Learning in Python. J. Mach. Learn. Res. 12, 2825–2830 (2011).

26. L’Yi, S., Wang, Q., Lekschas, F. & Gehlenborg, N. Gosling: A Grammar-based Toolkit for Scalable and Interactive Genomics Data Visualization. IEEE Trans. Vis. Comput. Graph. PP, (2021).

27. Nusrat, S., Harbig, T. & Gehlenborg, N. Tasks, Techniques, and Tools for Genomic Data Visualization. Comput. Graph. Forum 38, 781–805 (2019).

28. Gundersen, S. et al. Identifying elemental genomic track types and representing them uniformly. BMC Bioinformatics 12, 494 (2011).

29. Lieberman-Aiden, E. et al. Comprehensive mapping of long-range interactions reveals folding principles of the human genome. Science 326, 289–293 (2009).

30. Quinodoz, S. A. et al. Higher-Order Inter-chromosomal Hubs Shape 3D Genome Organization in the Nucleus. Cell 0, (2018).

31. Quinodoz, S. A. et al. SPRITE: a genome-wide method for mapping higher-order 3D interactions in the nucleus using combinatorial split-and-pool barcoding. Nat. Protoc. 17, 36–75 (2022).

32. Johnson, D. S., Mortazavi, A., Myers, R. M. & Wold, B. Genome-wide mapping of in vivo protein-DNA interactions. Science 316, 1497–1502 (2007).

33. Skene, P. J. & Henikoff, S. An efficient targeted nuclease strategy for high-resolution mapping of DNA binding sites. Elife 6, (2017).

34. Buenrostro, J. D., Giresi, P. G., Zaba, L. C., Chang, H. Y. & Greenleaf, W. J. Transposition of native chromatin for fast and sensitive epigenomic profiling of open chromatin, DNA-binding proteins and nucleosome position. Nat. Methods 10, 1213–1218 (2013).

35. Johnson, S. M., Tan, F. J., McCullough, H. L., Riordan, D. P. & Fire, A. Z. Flexibility and constraint in the nucleosome core landscape of Caenorhabditis elegans chromatin. Genome Res. 16, 1505–1516 (2006).

36. Core, L. J., Waterfall, J. J. & Lis, J. T. Nascent RNA sequencing reveals widespread pausing and divergent initiation at human promoters. Science 322, 1845–1848 (2008).

37. Hansen, R. S. et al. Sequencing newly replicated DNA reveals widespread plasticity in human replication timing. Proc. Natl. Acad. Sci. U. S. A. 107, 139–144 (2010).

38. Sheffield, N. C. & Bock, C. LOLA: enrichment analysis for genomic region sets and regulatory elements in R and Bioconductor. Bioinformatics 32, 587 (2016).

39. McInnes, L., Healy, J. & Melville, J. UMAP: Uniform Manifold Approximation and Projection for Dimension Reduction. arXiv [stat.ML] (2018).

40. Frankish, A. et al. GENCODE 2021. Nucleic Acids Res. 49, D916–D923 (2021).

41. Hsieh, T.-H. S. et al. Resolving the 3D Landscape of Transcription-Linked Mammalian Chromatin Folding. Mol. Cell (2020) doi:10.1016/j.molcel.2020.03.002.

42. Bonev, B. et al. Multiscale 3D Genome Rewiring during Mouse Neural Development. Cell 171, 557–572.e24 (2017).

43. Krietenstein, N. et al. Ultrastructural Details of Mammalian Chromosome Architecture. Mol. Cell 78, 554–565.e7 (2020).

44. Hsieh, T. H. S. et al. Mapping Nucleosome Resolution Chromosome Folding in Yeast by Micro-C. Cell (2015) doi:10.1016/j.cell.2015.05.048.

45. Banigan, E. J. et al. Transcription shapes 3D chromatin organization by interacting with loop-extruding cohesin complexes. bioRxiv 2022.01.07.475367 (2022) doi:10.1101/2022.01.07.475367.

46. Ladurner, R. et al. Sororin actively maintains sister chromatid cohesion. EMBO J. 35, 635–653 (2016).

47. Thiecke, M. J. et al. Cohesin-Dependent and -Independent Mechanisms Mediate Chromosomal Contacts between Promoters and Enhancers. Cell Rep. 32, 107929 (2020).

48. Kagey, M. H. et al. Mediator and cohesin connect gene expression and chromatin architecture. Nature 467, 430–435 (2010).

49. Mitter, M. et al. Conformation of sister chromatids in the replicated human genome. Nature 586, 139–144 (2020).

50. Davidson, I. F. et al. DNA loop extrusion by human cohesin. Science 366, 1338–1345 (2019).

51. Kim, Y., Shi, Z., Zhang, H., Finkelstein, I. J. & Yu, H. Human cohesin compacts DNA by loop extrusion. Science 366, 1345–1349 (2019).

52. Bannister, A. J. & Kouzarides, T. Regulation of chromatin by histone modifications. Cell Res. 21, 381–395 (2011).

53. Karmodiya, K., Krebs, A. R., Oulad-Abdelghani, M., Kimura, H. & Tora, L. H3K9 and H3K14 acetylation co-occur at many gene regulatory elements, while H3K14ac marks a subset of inactive inducible promoters in mouse embryonic stem cells. BMC Genomics 13, 424 (2012).

54. McLean, C. Y. et al. GREAT improves functional interpretation of cis-regulatory regions. Nat. Biotechnol. 28, 495–501 (2010).

55. Auerbach, R. K., Chen, B. & Butte, A. J. Relating genes to function: identifying enriched transcription factors using the ENCODE ChIP-Seq significance tool. Bioinformatics 29, 1922–1924 (2013).

56. Favorov, A. et al. Exploring Massive, Genome Scale Datasets with the GenometriCorr Package. PLoS Comput. Biol. 8, e1002529 (2012).

57. Yatskevich, S., Rhodes, J. & Nasmyth, K. Organization of Chromosomal DNA by SMC Complexes. Annu. Rev. Genet. 53, 445–482 (2019).

58. Cuadrado, A. & Losada, A. Specialized functions of cohesins STAG1 and STAG2 in 3D genome architecture. Curr. Opin. Genet. Dev. 61, 9–16 (2020).

59. Sumara, I., Vorlaufer, E., Gieffers, C., Peters, B. H. & Peters, J. M. Characterization of vertebrate cohesin complexes and their regulation in prophase. J. Cell Biol. 151, 749–762 (2000).

60. Losada, A., Yokochi, T., Kobayashi, R. & Hirano, T. Identification and characterization of SA/Scc3p subunits in the Xenopus and human cohesin complexes. J. Cell Biol. 150, 405–416 (2000).

61. Kojic, A. et al. Distinct roles of cohesin-SA1 and cohesin-SA2 in 3D chromosome organization. Nat. Struct. Mol. Biol. 25, 496–504 (2018).

62. Casa, V. et al. Redundant and specific roles of cohesin STAG subunits in chromatin looping and transcriptional control. Genome Res. 30, 515–527 (2020).

63. Wutz, G. et al. ESCO1 and CTCF enable formation of long chromatin loops by protecting cohesinstag1 from WAPL. Elife 9, (2020).

64. Rankin, S., Ayad, N. G. & Kirschner, M. W. Sororin, a Substrate of the Anaphase-Promoting Complex, Is Required for Sister Chromatid Cohesion in Vertebrates. Mol. Cell 18, 185–200 (2005).

65. Schmitz, J., Watrin, E., Lénárt, P., Mechtler, K. & Peters, J.-M. Sororin Is Required for Stable Binding of Cohesin to Chromatin and for Sister Chromatid Cohesion in Interphase. Curr. Biol. 17, 630–636 (2007).

66. Jalili, V. et al. The Galaxy platform for accessible, reproducible and collaborative biomedical analyses: 2020 update. Nucleic Acids Res. 48, W395–W402 (2020).

67. Younesy, H., Möller, T., Lorincz, M. C., Karimi, M. M. & Jones, S. J. . VisRseq: R-based visual framework for analysis of sequencing data. BMC Bioinformatics 16 Suppl 11, S2 (2015).

68. Kent, W. J., Zweig, A. S., Barber, G., Hinrichs, A. S. & Karolchik, D. BigWig and BigBed: enabling browsing of large distributed datasets. Bioinformatics 26, 2204–2207 (2010).

